# Conditional Protein Rescue (CPR) by Binding-Induced Protective Shielding

**DOI:** 10.1101/2020.06.16.154435

**Authors:** Andrew S. Gaynor, Wilfred Chen

## Abstract

An effective method to modulate the stability of proteins is essential to biological research. Herein, we describe a new technology that allows conditional stabilization of proteins based on masking of a degron tag by a specific intracellular protein cue. A target protein is fused to a degron tag and an affinity sensor domain. When the sensor detects its target protein, the degron is effectively concealed and the target protein is rescued. By introducing nanobodies as the sensor, we allow for virtually any endogenous protein to be targeted. In a model system using yeast cytosine deaminase, we demonstrate low cell death background yet maintain the ability to elicit strong activation and prodrug-mediated cell killing using GFP as the rescue protein. The flexibility in choosing different masking targets provides a straightforward method to generalize the strategy for conditional protein rescue in a wide range of biological contexts, including oncoprotein detection.

## Introduction

Conditional control of protein levels remains elusive for many biological applications. RNA interference (RNAi) destroys mRNA, but it can frequently be off-target or partially potent (Sigoillot and King, 2011). While small molecule-responsive transcriptional switches are frequently used to regulate mRNA levels, the overall dynamic is limited by the half-life of the target protein (Battle et al., 2015; Vogel and Marcotte, 2012; Wu et al., 2013). Another common method is the fusion of a degradation domain (DD) to a protein of interest (POI) (Li et al., 1998), which drastically reduces its half-life and allows faster fluctuations in the intracellular level (Mei et al., 2018; Sjaastad et al., 2018). As we recently reviewed (Chen et al., 2019), while several approaches can modulate protein degradation in response to a small molecule (Chung et al., 2015; Iwamoto et al., 2010; Lau et al., 2010), they do not allow protein concentration control in response to native cellular environments. Ideally, a modular platform that combines rapid protein turnover by DDs with temporal and autonomous responsiveness to cellular environments will greatly expand our ability to generalize the strategy for conditional protein rescue (CPR) in a wide range of biological contexts.

Coordinated degradation of cyclins is a key mechanism to ensure correct progression through the cell cycle (Harper et al., 2002; Morgan, 1997; Sherr and Roberts, 1999). This exquisite control between accumulation and depletion of cyclins is tightly regulated by changes in cellular protein information, suggesting a possible framework for CPR. One potential strategy is based on the Ac/N-End Rule pathway used for protein quality control, which recognizes and targets certain Nα-terminally acetylated resides for degradation (Oh et al., 2017; Shemorry et al., 2013; Zhang et al., 2010). Remarkably, the same acetylated residue is also necessary for proper interaction with cellular chaperones, which sterically shield the degradation domain and preserve properly folded proteins. The intriguing ability to shield the DD from initiating degradation has inspired the design of a new generation of artificial protein stability switches for conditional degradation. Insertion of a DD into the Jα-helix successfully shielded the DD-Jα-helix peptide within the LOV domain and arrested degradation. Irradiation with blue light unmasked the Jα-helix and restored degradation (Renicke et al., 2013). Similarly, a DD placed between two proteins was only activated upon release by protease cleavage (Jungbluth et al., 2010; Taxis et al., 2009). While these reports represent a first step towards CPR, they are unable to couple endogenous cellular cues to modulate degradation.

We sought to increase the practicality of CPR by using cellular protein cues to provide masking and unmasking of DDs. In this design, a small sensor domain is appended to the DD. When a binding target is present, the DD is effectively concealed, and the target protein is rescued (Fig. 1A). We demonstrated that effective CPR can be executed using both covalent SpyTag/SpyCatcher conjugation and non-covalent nanobody/antigen interaction. Selective rescue of the yeast cytosine deaminase enabled strong prodrug activation and targeted cell killing.

**Figure 1.**
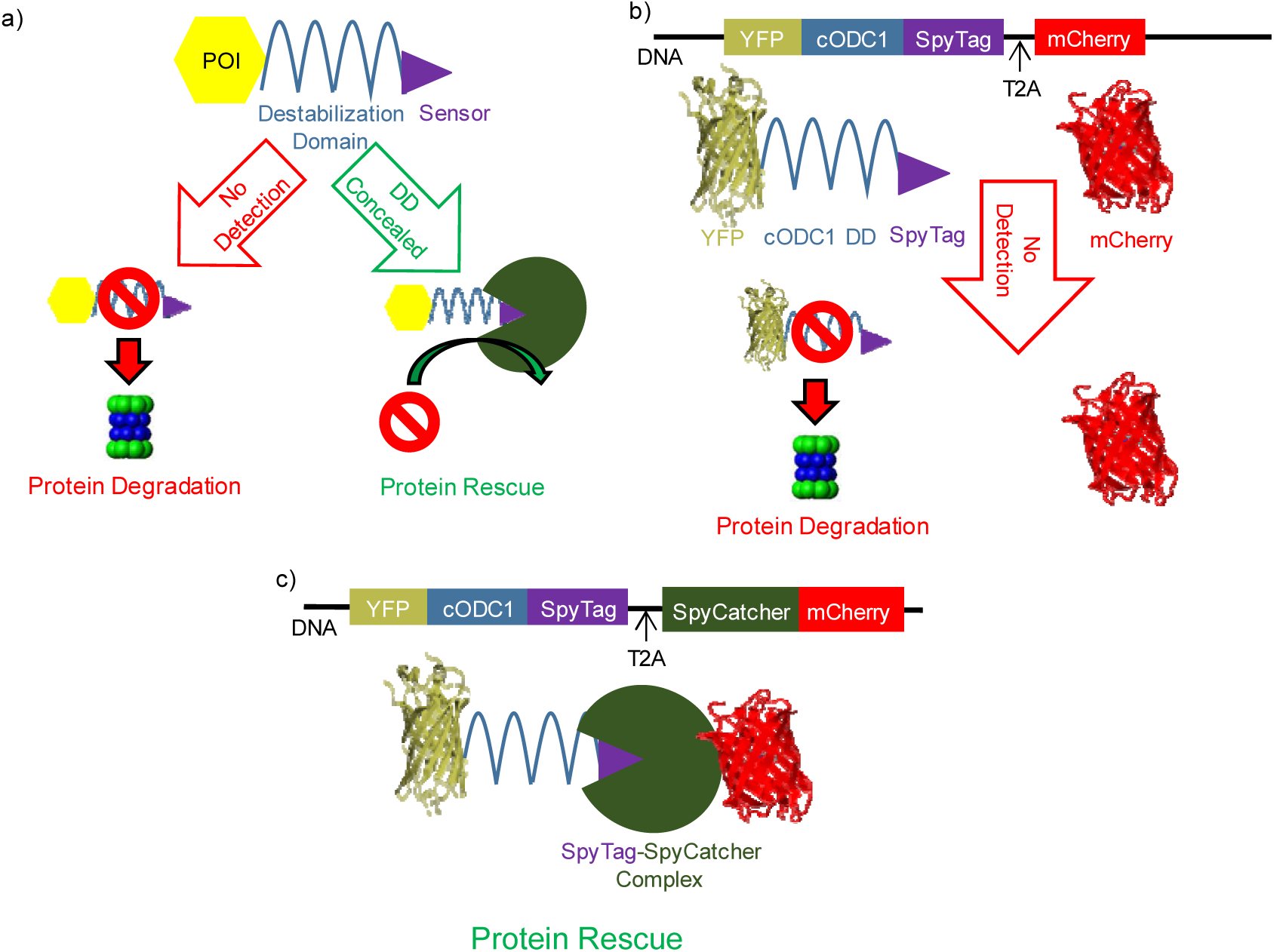
Conditional protein rescue (CPR) via masking the DD. a) The DD (blue squiggle) contains a small sensor domain (purple triangle) fused to its C-terminus. In the absence of the corresponding binding target to the sensor, the POI (yellow hexagon) is recruited to the proteasome via DD interaction (red symbol) and degradation proceeds (left). Interaction with the target (green cut-out circle) conceals the DD from the proteasomal recruitment, and the POI is rescued from degradation (right). b) YFP is fused to the cODC1 DD and SpyTag (sensor) and co-expressed with mCherry as a transfection marker. YFP is degraded by proteasome recognition of cODC1, and mcherry remains. c) When mcherry is fused to the SpyCatcher (target), the SpyTag sensor recruits SpyCatcher-miRFP670, sterically concealing cODC1 and rescuing YFP by CPR.

## Results

### Conditional protein rescue by covalent SpyTag/SpyCatcher conjugation

To evaluate the feasibility of CPR, we first utilized the SpyCatcher and SpyTag system, which provides the most stable *in vivo* interaction because of covalent conjugation (Zakeri et al., 2012). A well-characterized synthetic cODC1-like C-degron tag was used as an effective DD with kinetics that allow for rescue to occur (Renicke et al., 2013). By fusing the DD-SpyTag to a fluorescent reporter, we generated YFP-cODC1-SpyTag, an unstable complex that can be rescued by SpyCatcher. We employed mCherry as an orthogonal transfection reporter (Fig. 1B). Both the YFP fusion and mCherry were expressed under one promotor by use of a polycistronic viral T2A self-cleaving sequence (Holst et al., 2006; Szymczak et al., 2004). To induce rescue of YFP, SpyCatcher was fused to mCherry for easy tracking (Fig. 1C).

To evaluate the rescue efficiency, HeLa cells were transfected with SpyCatcher-mCherry:T2A:YFP-cODC1-SpyTag or the control, mCherry:T2A:YFP-cODC1-SpyTag, without SpyCatcher. Expression of both proteins was tracked by fluorescent microcopy and western blot over 60 h. Western blot analysis (Fig. 2A) demonstrated that mCherry was detected consistently in both constructs roughly 15 h post-transfection. YFP gradually disappeared in cells expressing only mCherry, while a strong band corresponding only to the ligated YFP products was detected for cells expressing SpyCatcher-mCherry. The absence of any un-ligated YFP with SpyCatcher-mCherry co-expression highlights that ligation between SpyTag and SpyCatcher is solely responsible for YFP rescue due to shielding of the DD (Fig. 2A, top right box). This is further supported by the fluorescent images (Supplementary Fig. 1) demonstrating efficient YFP rescue due to DD shielding by SpyTag-SpyCatcher ligation.

**Figure 2.**
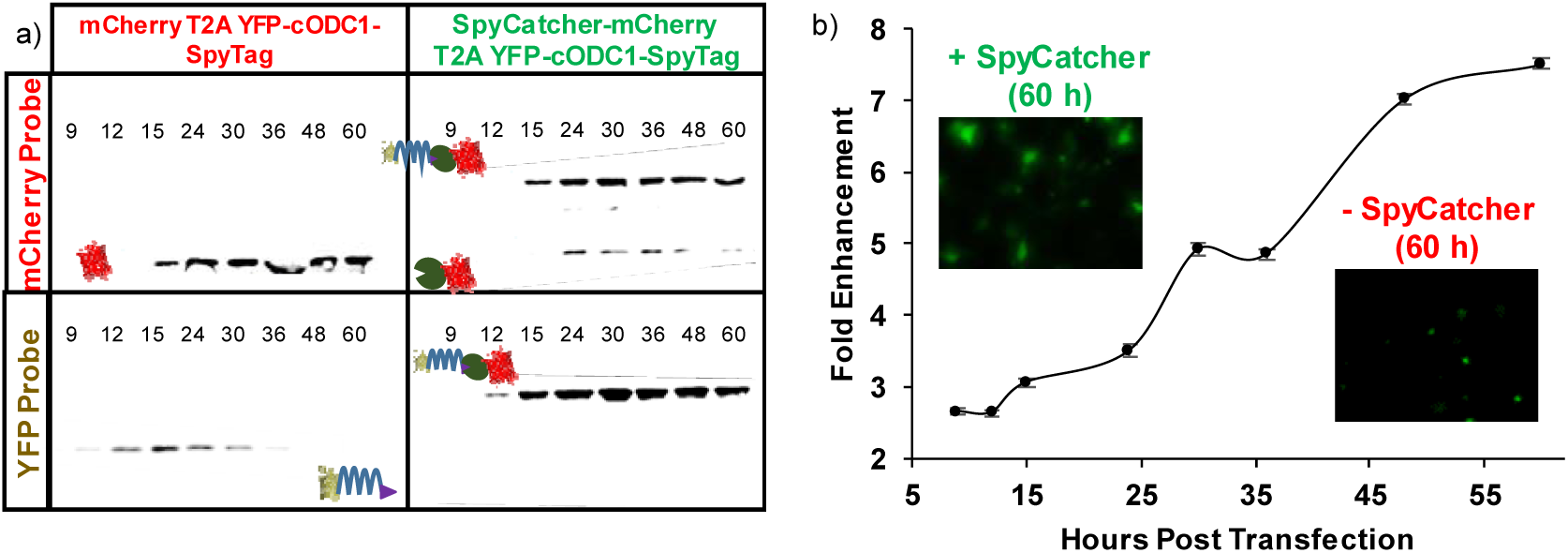
YFP rescue from cODC1-mediated degradation via SpyTag-SpyCatcher interaction. a) Western blotting of HeLa cell lysate. Expression of YFP and mCherry/mCherry-SpyCatcher were by their respective antibodies. The upward shift in the protein size for the mcherry-SpyCatcher samples was the result of SpyTag-SpyCatcher conjugation. b) Flow cytometry quantification of YFP enhancement by CPR. miRFP670, a near-infrared fluorescent protein with a completely orthogonal signal to YFP on the flow cytometer, was used in place of mcherry. Fold enhancement is YFP signal normalized to miRFP670 expression in the SpyCatcher-miRFP670 fusion sample relative to the control with no SpyCatcher expression. Error bars represent 95% confidence intervals.

To quantify CPR more accurately, miRFP670 — a near-infrared, monomeric, fluorescent protein with a completely orthogonal signal to YFP on the flow cytometer (Shcherbakova et al., 2016) — was fused similarly to mCherry to generate SpyCatcher-miRFP670:T2A:YFP-cODC1-SpyTag and the control, miRFP670:T2A:YFP-cODC1-SpyTag. Flow cytometry showed that CPR enhancement increased throughout the entire time course, with roughly 7.5-fold increase in the YFP signal after 60 h (Fig. 2B). Western blots confirmed a similar size increase as a result of the covalent conjugation between SpyCatcher and SpyTag (Supplementary Fig. 2).

### Use of non-covalent interactions for CPR

Although CRP was correctly executed using covalent conjugation between SpyTag and SpyCatcher, most intracellular interactions are non-covalent in nature. To show that non-covalent interaction can also be used to provide similar shielding effects, we replaced the SpyTag/SpyCatcher pair with the well-known Src homology 3 (SH3) domain and its corresponding binding ligand, wLig (k_D_ = 10 μM) (Dueber et al., 2007; Li, 2005). A similar shielding effect was observed albeit at reduced efficiencies, confirming that even a weak non-covalent interaction is sufficient to provide adequate masking of the DD (Supplementary Fig. 3). Again, no rescue was observed when the SH3 domain is absent, highlighting again the importance of specific interaction for proper DD masking (Supplementary Fig. 3).

In order to adapt this technology towards more relevant cellular targets, a small, monomeric sensor capable of interacting with endogenous proteins with high specificity is required. Camel single-domain antibody fragments, or nanobodies, are ideal because of their relative small size (∼13kDa) and the ability to generate high-affinity nanobodies for virtually any protein target (Kubala et al., 2010; Saerens et al., 2005). To investigate whether the degradation phenotype could be preserved even after addition of a nanobody near the DD, an anti-GFP nanobody (GBP1, k_D_ ∼1 nM) was first fused to the C-terminus of an miRFP670-cODC1 fusion (Fig. 3A). Unlike conjugation of a SpyCatcher-fusion onto an adjacent SpyTag to cOCD1, no masking of the DD was observed as virtually no miRFP670 signal was detected (Supplementary Fig. 4, left side). This is somewhat unexpected as a small structural nanobody was physically tethered next to the DD. We speculate that the steric masking of the DD may be size dependent. To test this hypothesis, we fused a larger maltose-binding protein (MBP; 43 kDa) to the C-terminus of GBP1. This resulted in improved miRFP670 signal (Supplementary Fig. 4, right side), an outcome consistent with the proposed enhanced DD masking and miRFP670 rescue.

**Figure 3.**
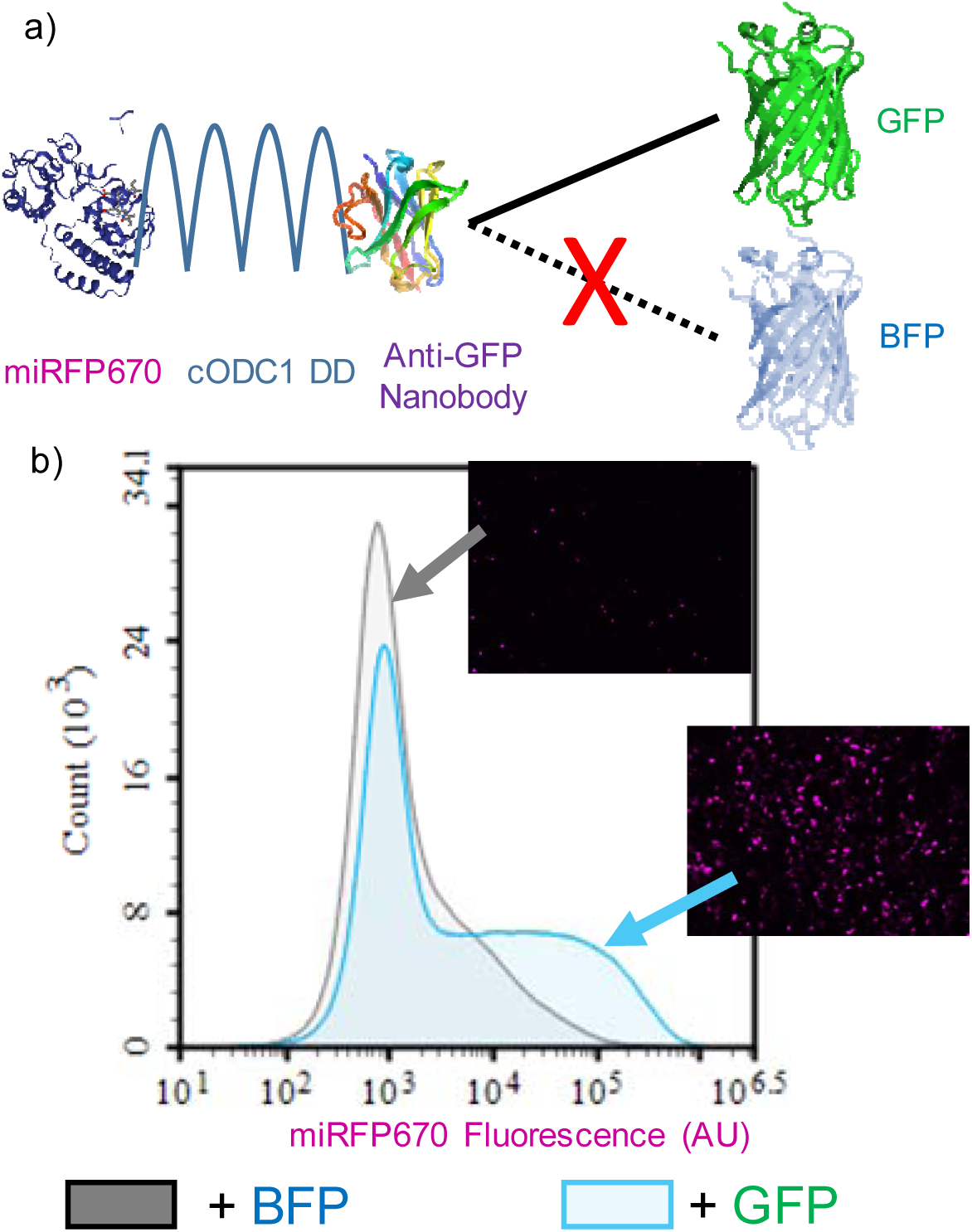
Rescuing a POI using non-covalent nanobody-antigen interactions. a) miRFP670 is fused to the cODC1 DD and an anti-GFP nanobody (GBP1), which still maintains its inherently unstable feature. HeLa cells expressing miRFP670-cODC1-GBP1 were co-transfected with either BFP or GFP for CPR. Co-expression with BFP alone did not result in miRFP670 rescue due to a lack of interaction with GBP1, while co-expression with GFP restored miRFP670 signal due to DD masking. b) Flow cytometry quantification of miRFP670 fluorescence in the presence of BFP (grey) and GFP (blue) after 48 h. Inserts show fluorescent microscopy images of miRFP670 of each sample.

After establishing that a small nanobody GBP1 can be fused after the DD without impacting degradation, we next investigated whether protein rescue could be attained based on GBP1 and GFP interaction. In the presence of BFP, which could not associate with GBP1, miRFP670 was still efficiently degraded. In contrast, expression of GFP efficiency rescued miRFP670 from degradation due to GFP shielding of the DD (Fig. 3B and Supplementary Fig. 4). Somewhat surprisingly, GFP failed to induce as effective CPR when we used GBP6, a nanobody that binds GFP at a different epitope than GBP1 (Tang et al., 2013), suggesting that interacting orientation, in addition to the size of the rescuing protein, is also important for CPR (Supplementary Fig. 5). These results provide the feasibility to repurpose nanobody-antigen interactions to elicit CPR for many different synthetic biology applications of practical interest.

### Engineering CPR for prodrug activation

One of the most pressing needs in cancer treatment is to distinguish cancer versus healthy cells. Prodrug targeting offers a layer of therapeutic control due to the innocuous nature of prodrugs. Yeast cytosine deaminase (yCD) is a prodrug-converting enzyme (PCE) that transforms the innocuous 5-fluorocytosine (5-FC) into the cytotoxic 5-fluorouracil (5-FU), and it has been used successfully for the treatment of glioblastoma (Polak et al., 1976; Zhang et al., 2014). Previously, we demonstrated the ability to regulate yCD activity using a small molecule-dependent rescue system, but this approach lacked any autonomous ability to distinguish cancer cells from healthy cells (Gaynor and Chen, 2017). To adapt CPR for prodrug targeting, yCD was used as the POI to test how well this strategy can control 5-FC activation. GFP again served as a visually trackable surrogate for a cancer-relevant protein. The ability to trigger cell death by 5-FU was used to indicate the overall efficiency of the conditional PCE therapy. A dye that only crosses the leaky cell membrane of dead cells was used as a visible indicator of cell viability. As expected, 5-FU killed large quantities of cells regardless of GFP, while cell viability was high when no drugs were administered (Fig. 4). When treated with 5-FC, only cells co-expressing GFP were killed in similar quantities to those being treated directly with 5-FU (Fig. 4). Although the degree of cell killing in the absence of GFP but with 5-FC is slightly higher than cells without 5-FC addition (Fig. 4B), this undesired outcome can be rectified by using a stronger degradation signal (*e.g.* UbL) or a combination of multiple DDs.

**Figure 4.**
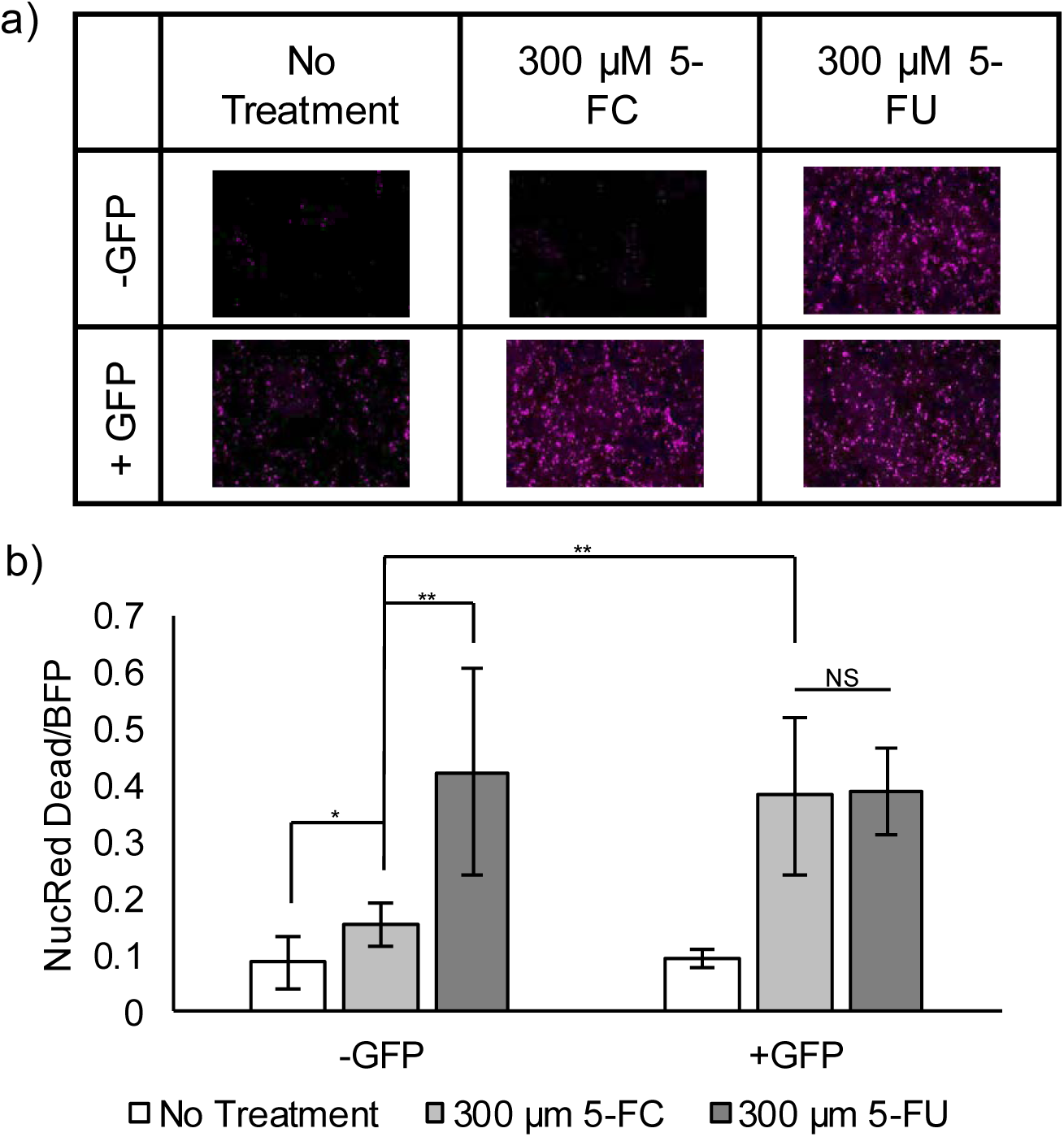
Controlling yCD activity via protein-nanobody interaction-mediated rescue. a) Fluorescent images of a cell death dye. Presence of the dye (pink) indicates a dead cell. Cells die in large numbers in the presence of 5-FU. Cells are killed by 5-FC, the prodrug, only when GFP rescues yCD by stabilizing the DD-nanobody fusion. b) Quantification of all fluorescent images, normalizing NucRed Dead dye to BFP, the protein transfection marker. Cells were transfected and either treated with no drugs (No Treatment), 5-FC, or 5-FU (*n* = 10; * = *p* < 0.05; ** = *p* < 0.01; NS = no statistical significant difference).

### Tuning CPR by using a stronger proteasome binding motif

We next sought to improve the design to eliminate the background further. Previously, it has been reported that an unstructured domain and a proteasomal targeting moiety are both necessary for efficient proteasomal degradation (Prakash et al., 2009). To determine if our CPR design could block access to the unstructured cOCD1 domain in the presence of a second proteasomal targeting moiety, we fused one copy of the ubiquitin-like (UbL) domain to the N-terminus of YFP (Stack et al., 2000). UbL is derived from the Rad23 protein and has been shown to target its fusion partners directly to the proteasome more effectively than the cODC1 tag (Elsasser et al., 2002; Yu et al., 2016). Fusing a UbL domain to the N-terminus of YFP enhanced the overall degradation, demonstrating that CPR can be tuned to achieve varying activation levels and signal to background ratios (compare the disappearance of YFP bands in Fig. 5B with Fig. 2A). Neither fluorescent microscopy (Fig. 5A and Supplementary Fig. 6) nor western blot (Fig. 5B) could detect YFP in the absence of SpyCatcher-mCherry. Co-expression of SpyCatcher-mCherry was again able to rescue YFP, although the rescued YFP level was lower than without the UbL domain (Fig. 5A). The increase in protein degradation kinetics competes more aggressively with the SpyTag-SpyCatcher reaction, resulting in less rescue. This result highlights the modularity of our approach in adjusting signal background and rescue intensity and its ability to conceal unstructured domains from a proteasome in a UbL-tagged target.

**Figure 5.**
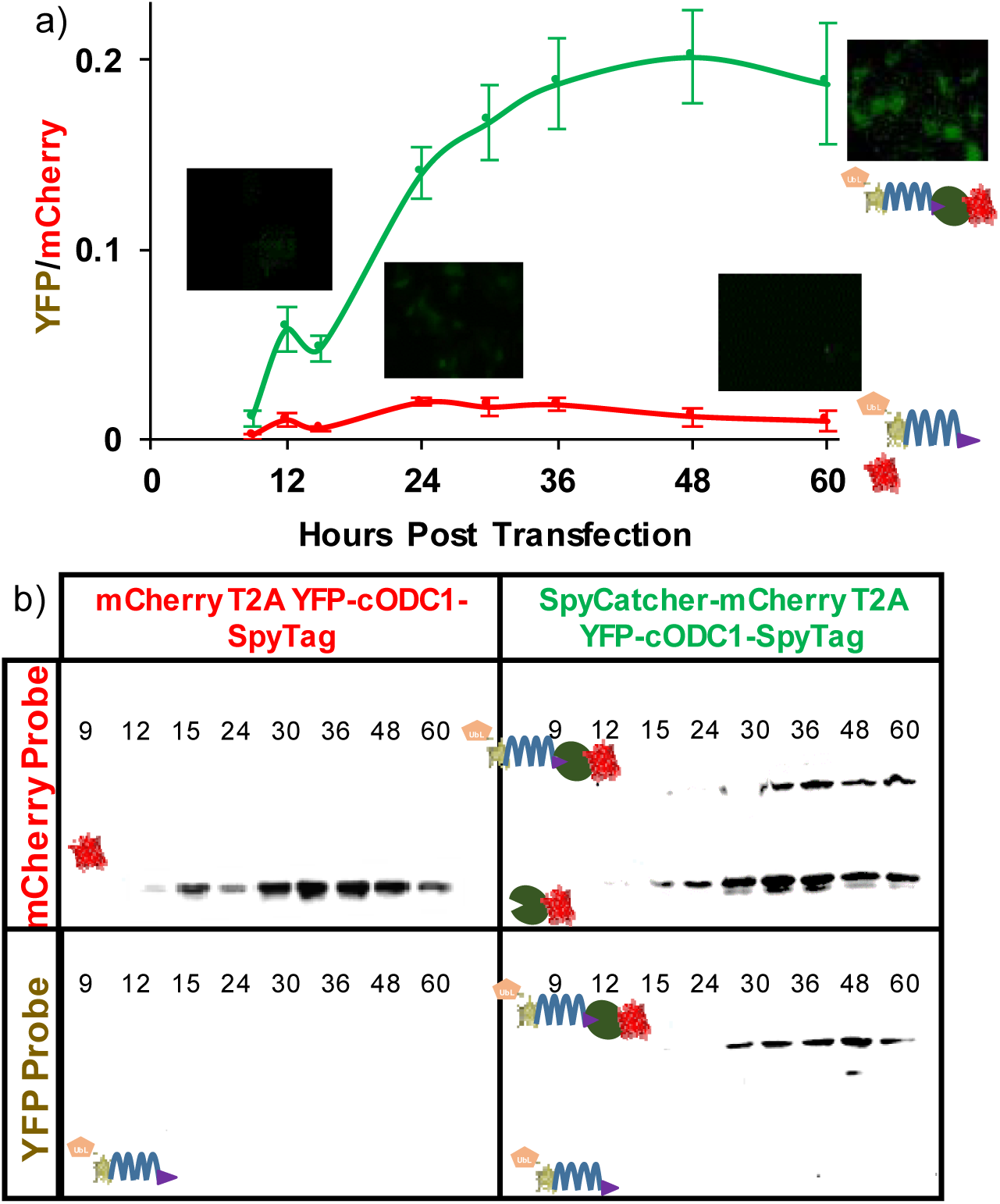
Tuning YFP rescue using the stronger proteasome binding UbL domain to improve degradation kinetics. a) Quantification of fluorescent microscopy measuring YFP intensity normalized by mCherry intensity. Compared to designs without UbL (see Fig. 2), background YFP intensity was decreased (red line), but the ability of YFP to be rescued decreased as well (green line). The images show HeLa cells with the YFP signal (green) 9 h and 60 h post transfection. Error bars represent ± 95% confidence interval (n = 5). b) Western blotting of HeLa cell lysate. The UbL domain is effective in eliminating any detectable traces of YFP expression without rescue (lower left box). Co-expression with SpyCatcher rescued YFP from degradation (lower right box).

### CPR for N-End Rule Degrons

Encouraged by the CPR results using the cOCD1 C-degron, we next turned our attention to the N-end rule protein degradation pathway. Because the N-end rule substrates are recognized by specific binding proteins known as N-recognins, which deliver these substrates to the 26S proteasome for destruction (Choi et al., 2010; Matta-Camacho et al., 2010), chaperones are able to protect their targets via steric interference (Zhang et al., 2010). We reasoned that expressing a sensing nanobody directly following a destabilizing N-terminus residue as a fusion to a POI should result in rescue when the corresponding nanobody’s target is co-expressed.

To conduct CPR using an N-end rule degron, we relied upon the ubiquitin (Ub) fusion technique, in which Ub is added to the N-terminus of a POI. The Ub domain is subsequently cleaved by an endogenous deubiquitylase, exposing the desired N-terminus residue for destabilization (Bachmair et al., 1986). Using this strategy, we generated three Ub:X-GBP1-miRFP670 fusions, where X is the resulting N-terminal residue: methionine (M, half-life = 30 hr), leucine (L, half-life = 5.5 hr), or arginine (R, half-life = 1.0 hr) (Gonda et al., 1989). These constructs were co-expressed with either BFP or GFP. Co-expression of BFP resulted in weak miRFP670 fluorescence scaling to the reported half-lives of the N-terminus residues tested. In contrast, co-expressing with GFP resulted in a dramatic rescue of miRFP670, showing more fluorescence than rescuing with the cODC1 degron (Fig. 6a and Supplementary Fig. 7). Due to its extremely low background yet high level of rescue, an Arg N-terminus degron elicited an unprecedented >50-fold increase in protein fluorescence. To ensure that CPR was not a phenomenon specific to GBP1 and GFP-mediated rescue, we replaced GBP1 with LaM4, a nanobody that detects mCherry (Fridy et al., 2014), to create EGFP-cODC1-LaM4 and three different Ub:X-LaM4-EGFP fusions. For all constructs, only co-expression with mCherry resulted in higher EGFP fluorescence, and the N-end rule CPR outperformed C-end rule (Supplementary Fig. 8).

**Figure 6.**
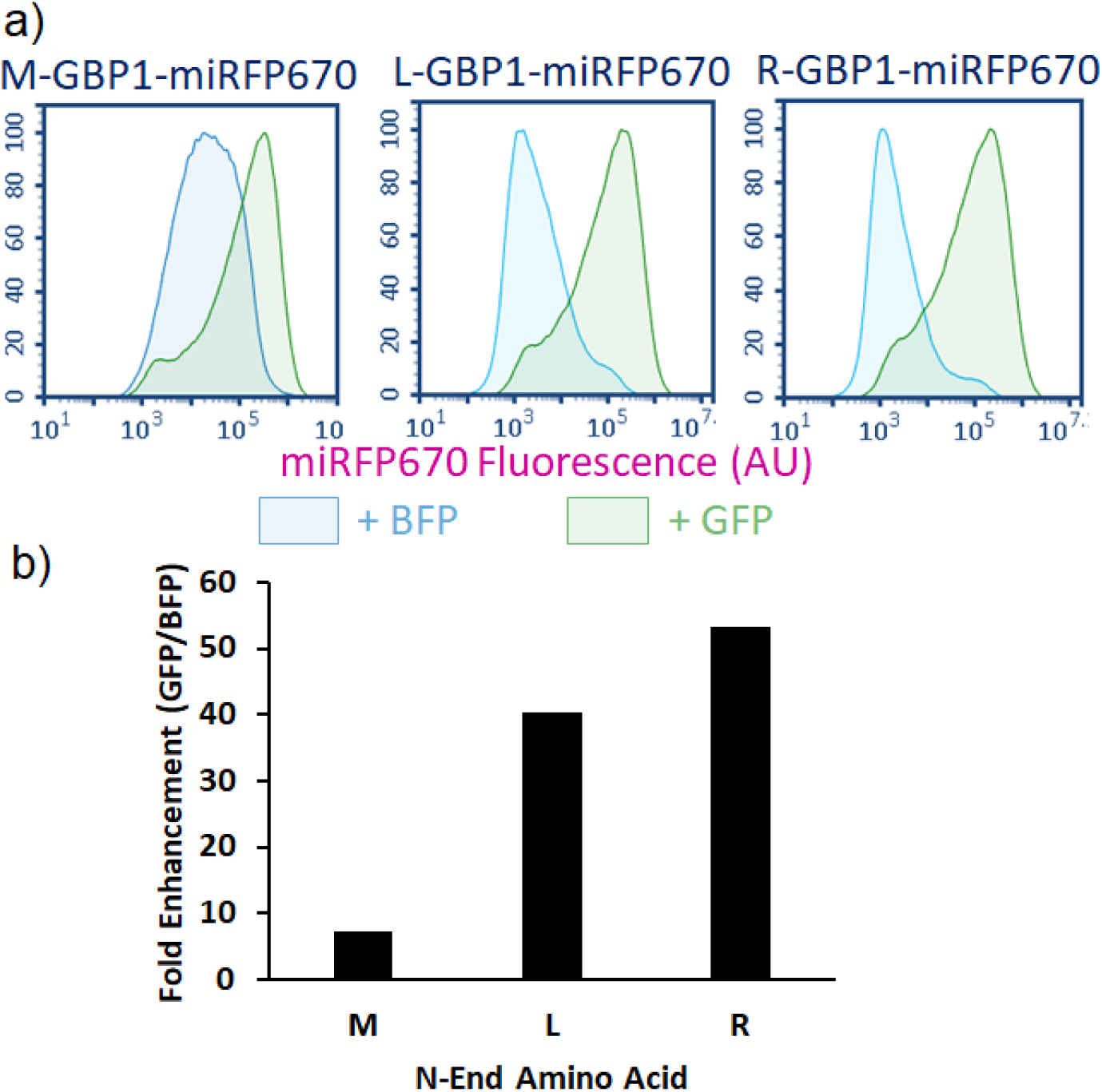
Rescuing protein by blocking N-end rule-mediated degradation. a) When BFP is co-expressed, miRFP670 rescue does not occur, and the residual fluorescence levels scale well with the reported half-lives of proteins with the respective N-terminal amino acids. However, co-expression of GFP resulted in a significant increase in miRFP670 fluorescence levels. Furthermore, fluorescence levels of rescued protein are comparable regardless of which N-terminal amino acid is used. b) The fold enhancement measured for each N-terminal amino acid is plotted as a function of the median miRFP670 fluorescence when co-expressed with GFP divided by the median when co-expressed with BFP (background fluorescence). Each N-terminal amino acid noted some enhancement, with Arg measuring more than 50x enhancement. Error bars represent a 95% confidence interval.

### CPR for the Detection of HPV-Positive Cells

To illustrate the broader applicability of our CPR approach toward native protein targets, we next extended our design to detect human papillomavirus (HPV)-positive cells. HPV is a known oncovirus that mainly relies on two proteins to induce carcinogenesis in cervical cells: E6, a major suppressor of apoptosis, and E7, a driver of the cell cycle (Jansma et al., 2014; Moody and Laimins, 2010; Senba and Mori, 2012). Using E7 as a HPV marker, we exploited nE7, a nanobody that detects E7 (Li et al., 2019), to generate Ub:R-nE7-mCherry to execute CPR. We transfected this construct and the control Ub:R-GBP1-mCherry into both HPV-positive HeLa cells and HPV-negative HEK293T cells. The HEK293T cells showed similar low levels of mCherry fluorescence regardless of which nanobody was used to perform CPR (Fig. 7). However, while GBP1 resulted in low levels of mCherry in the HeLa cells, nE7 resulted in a roughly 3-fold increase in mCherry, demonstrating that CPR is a powerful technique for detecting even low cellular levels of a cellular target protein.

**Figure 7.**
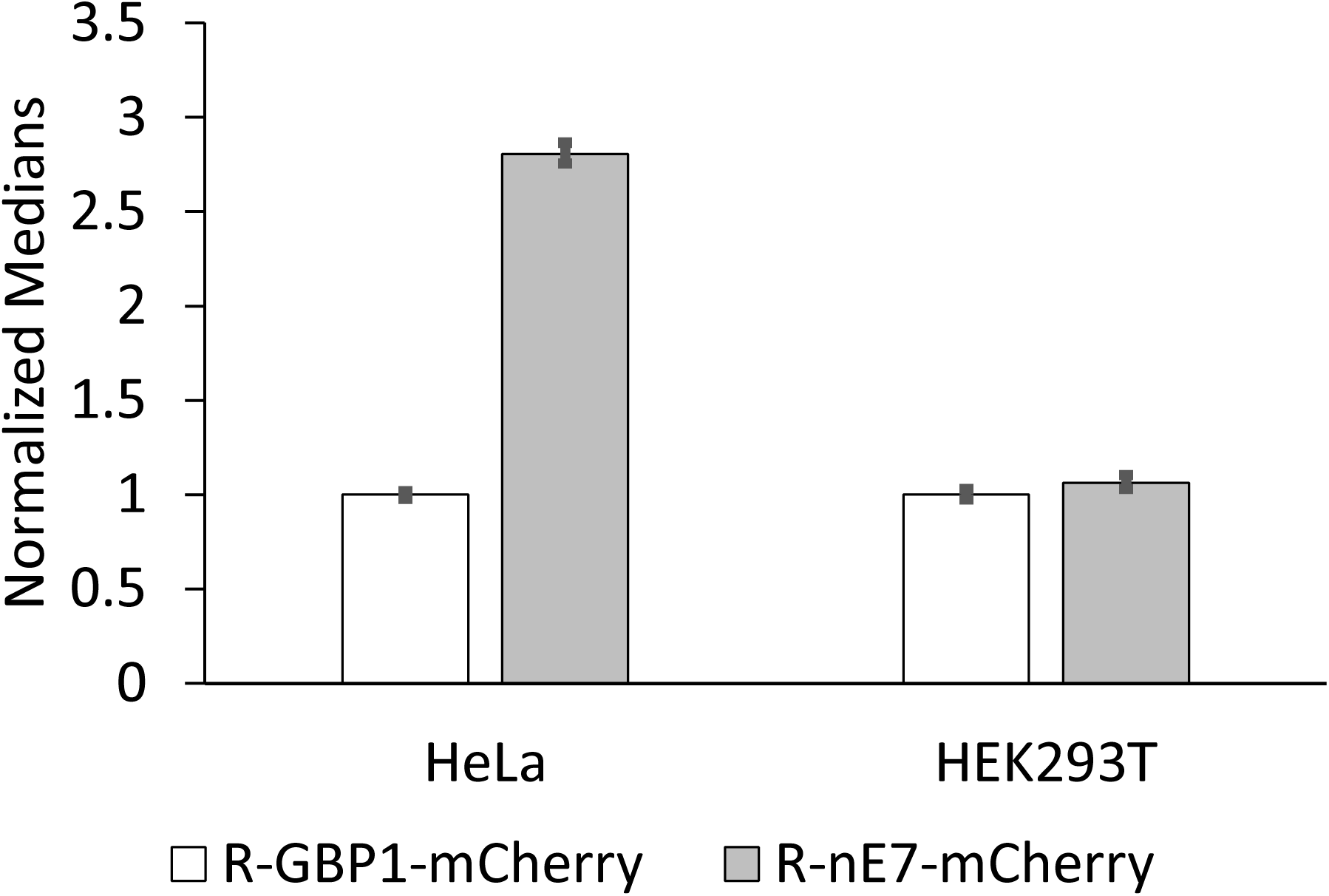
Detecting an endogenous cancer marker using CPR. HeLa cells are cancerous as a consequence of infection with HPV. These viral proteins provide a specific marker for HeLa cells that can be detected by nE7 nanobody (left). HEK293T cells do not contain this marker, and therefore no statistically significant difference is observed. For both cell types, median fluorescence is normalized to R-GBP1-mCherry fluorescence (background). Error bars represent 95% confidence intervals.

## Discussion

We report here a new synthetic biology framework to elicit CPR based on proteomic information. To our knowledge, this is the first report that allows for the rescue of a target protein from degradation using a second protein as a masking agent. Although the initial feasibility was demonstrated using the SpyTag/SpyCatcher bioconjugation pair, even non-covalent interactions can be used to achieve similar rescue efficiencies. The modularity of the design allows the addition of a UbL proteasome-targeting domain to eliminate background while still allowing rescue of a POI. The use of nanobodies as a small sensing domain removes the limit on the potential target pool and creates a new synthetic biology framework by allowing endogenous cellular proteins to decide the fate of a POI. We demonstrated this feasibility by detecting E7, a protein unique to HPV-positive cells. The availability of DDs with a wide range of degradation kinetics, including the N-end rule, offers the possibility to elicit rescue by an endogenous protein in a threshold-dependent manner. By combining different DDs and sensing domains, it may be possible to generate more complex, multi-input protein logic gates to help further differentiate between disease and healthy cells for therapeutic applications.

## Materials and Methods

### Plasmid construction

All constructs were prepared using standard molecular cloning techniques and cloned into pcDNA3.1(+) (Invitrogen). All oligonucleotides were ordered from Integrated DNA Technology (Coralville, IA) and purified via standard desalting. All enzymes were purchased from New England Biolabs (Ipswich, IA) and used per the manufacturer’s protocol with the provided buffers. All overlapping oligos were first 5’ phosphorylated with T4 polynucleotide kinase (PNK) treatment, and then were heat denatured and slow cooled to allow for proper hybridization before ligation.

### mCherry:T2A:YFP-cODC1-SpyTag

YFP was PCR amplified and double digested with *AflII* and *XhoI*. The DNA sequences for oCDC1-SpyTag were ordered as overlapping oligonucleotides as ultrameres with appropriate overhangs to make them complimentary to *XhoI* and *ApaI*. The vector pcDNA3.1(+) was double digested with *AflII* and *ApaI* to generate the backbone, and YFP and cODC1-SpyTag were ligated simultaneously using T4 DNA Ligase per the manufacturer’s protocol to generate YFP-cODC1-SpyTag. Finally, mCherry was PCR amplified with a reverse primer that included the T2A region in the non-overlapping region, and this product was double digested with *NheI* and *AflII*. YFP-cODC1-SpyTag was double digested with *NheI* and *AflII*, and mCherry:T2A was ligated, generated mCherry:T2A:YFP-cODC1-SpyTag.

### SpyCatcher-mCherry T2A:YFP-cODC1-SpyTag

SpyCatcher was codon optimized and ordered as a gBlock gene fragment. SpyCatcher was then PCR amplified and double digested with *NheI* and *EcoRI*. mCherry:T2A was PCR amplified with the same reverse primer as above, but the forward primer provided an N-terminal *EcoRI* site, and this product was double digested with *EcoRI* and *AflII*. mCherry:T2A:YFP-cODC1-SpyTag was double digested with *NheI* and *AflII* to remove mCherry:T2A and generate the backbone into which SpyCatcher and mCherry:T2A were simultaneously ligated, generating SpyCatcher-mCherry:T2A:YFP-cODC1-SpyTag.

### miRFP670 constructs

miRFP670 was PCR amplified and double digested with *NheI* and *HindIII*. The T2A polycistronic site was ordered as two overlapping oligonucleotides with overhangs to provide *HindIII* and *AflII* complimentary sites. mCherry:T2A:YFP-cODC1-SpyTag was double digested with *NheI* and *AflII* to remove mCherry:T2A. miRFP670 and T2A were simultaneously ligated with the backbone to generate miRFP670:T2A:YFP-cODC1-SpyTag. Next, miRFP670 was PCR amplified with overhangs providing *EcoRI* and *HindIII* sites, and the product was double digested at those sites. The human codon optimized SpyCatcher was PCR amplified and double digested with *NheI* and *EcoRI* as described above. miRFP670:T2A:YFP-cODC1-SpyTag was double digested with *NheI* and *HindIII* to remove miRFP670, and SpyCatcher and miRFP670 was simultaneously ligated into the backbone, generating SpyCatcher-miRFP670:T2A:YFP-cODC1-SpyTag. Finally, SH3 was PCR amplified. SH3 and SpyCatcher-miRFP670:T2A:YFP-cODC1-SpyTag were double digested with *NheI* and *EcoRI* to remove SpyCatcher, and SH3 was ligated to generate SH3-miRFP670:T2A:YFP-cODC1-SpyTa*SH3* and *SH3Lig constructs*. SH3 was PCR amplified and double digested with *NheI* and *EcoRI*. SH3Lig was ordered as a pair of overlapping oligonucleotides with overhangs providing for *XbaI* and *ApaI* complementation sites. Previous plasmids could be double digested with *NheI* and *EcoRI* (to install SH3) or *XbaI* and *ApaI* (to install SH3Lig) to generate SpyCatcher-mCherry:T2A:YFP-cODC1-SH3Lig, SH3-mCherry:T2A:YFP-cODC1-SpyTag, or SH3-mCherry:T2A:YFP-cODC1-SH3Lig.

### miRFP670-cODC1-SpyTag-GBP1

SpyTag was ordered as a pair of overlapping oligonucleotides providing overhangs with *XbaI* and *BamHI*. The GFP Nanobody (GBP1), *aka* GFP Binding Protein 1 (GBP1), was PCR amplified and double digested with *BamHI* and *ApaI*. mCherry:T2A:YFP-cODC1-SpyTag was double digested with *XbaI ApaI* to remove SpyTag, and SpyTag and GBP1 were simultaneously ligated into the backbone generating mCherry:T2A:YFP-cODC1-SpyTag-GBP1. Next, miRFP670 was PCR amplified and double digested with *AflII* and *XhoI*,and mCherry:T2A:YFP-cODC1-SpyTag-GBP1 was double digested with *XhoI* and *ApaI* in order to purify cODC1-SpyTag-GBP1. pcDNA3.1(+) was double digested with *AflII* and *ApaI*, and miRFP670 and cODC1-SpyTag-GBP1 were simultaneously ligated into the vector to generate miRFP670-cODC1-SpyTag-GBP1.

### yCD constructs

BFP was PCR amplified and double digested with *NheI* and *ClaI*. A T2A site was ordered as overlapping ultramers with overhangs providing *ClaI* and *AflII* complementation sites. mCherry:T2A:YFP-cODC1-SpyTag-GBP1 was double digested with *NheI* and *AflII* to remove mCherry:T2A. BFP and T2A were simultaneously ligated into the cut vector to generate BFP:T2A:YFP-cODC1-SpyTag-GBP1. A second BFP was PCR amplified and double digested with *EcoRI* and *ClaI*. SH3 was again PCR amplified similar to above and double digested with *NheI* and *EcoRI*. BFP:T2A:YFP-cODC1-SpyTag-GBP1 was then double digested with *NheI* and *ClaI*, and SH3 and BFP were ligated to generate SH3-BFP:T2A:YFP-cODC1-SpyTag-GBP1. yCD was PCR amplified and double digested with *AflII* and *Xho*I. SH3-BFP:T2A:YFP-cODC1-SpyTag-GBP1 was also double digested with *AflII* and *XhoI* to remove YFP, and yCD was ligated in its place generating SH3-BFP:T2A:YFP-cODC1-SpyTag-GBP1. To generate the rescuing construct, EGFP was PCR amplified and double digested with *NheI* and *EcoRI*. The previous construct was double digested with the same enzymes to remove SH3, and GFP was ligated in its place generating GFP-BFP:T2A:YFP-cODC1-SpyTag-GBP1.

### UbL constructs

A single copy of the UbL domain was PCR amplified and double digested with *AflII* and *ClaI*. YFP was also PCR amplified and double digested with *ClaI* and *XhoI*. Both mCherry:T2A:YFP-cODC1-SpyTag and SpyCatcher-mCherry:T2A:YFP-cODC1-SpyTag were double digested with *AflII* and *XhoI* to remove YFP, and UbL and YFP were simultaneously ligated into the cut vector to generate mCherry:T2A:UbL-YFP-cODC1-SpyTag and SpyCatcher-mCherry:T2A:UbL-YFP-cODC1-SpyTag, respectively.

### N-end rule constructs

Ub-R-GFP was a gift from Nico Dantuma (Addgene plasmid # 11939; http://n2t.net/addgene:11939; RRID:Addgene_11939). Site directed mutagenesis was performed to generated Ub M-GFP and Ub-L-GFP using Q5 Hot Start High-Fidelity Polymerase New England Biolabs (Ipswich, IA) according to the manufacture’s protocol. KpnI and BamHI restriction sites were introduced via mutagenesis between the N-terminal amino acid and GFP, and GBP1 was inserted into these sites. Finally, GFP was excised using BamHI and NotI, and miRFP670 was ligated in its place to generate Ub X-GBP1-miRFP670.

### LaM4 constructs

To generate EGFP-cODC1-LaM4, EGFP was PCR amplified to include AflII and XhoI restriction sites. miRFP670-cODC1-GBP1 was digested with AflII and XhoI, and EGFP was ligated to generate EGFP-cODC1-GBP1. GBP1 was excised using BamHI and ApaI, and LaM4 was ligated in its place. N-End rule constructs were generated by excising GBP1 from Ub X-GBP1-EGFP using KpnI and BamHI; LaM4 was ordered as a gene fragment and PCR amplified to include the same restriction sites, and the ligation product yielded Ub X-LaM4-EGFP.

### nE7 constructs

Ub R-GBP1-miRFP670 was digested with KpnI and BamHI. The nE7 nanobody was ordered as a gene fragment from IDT, PCR amplified to include KpnI and BamHI restriction sites, and ligated into the vector. This subclone was subsequently digested with AgeI and NotI, and mCherry was PCR amplified and cloned into place to yield Ub R-nE7-mCherry. To generate the control construct, Ub R-GBP1-miRFP670 was digested with AgeI and NotI, and mCherry was PCR amplified and ligated into the vector to yield Ub R-GBP1-mCherry.

### Cell culture

HeLa cells were maintained in T150 tissue culture flasks (Thermo Fisher) in complete media, *i.e.* Minimum Essential Media (MEM, Cellgro) supplemented with 10% fetal bovine serum (FBS, Corning), 10 U mL^-1^ penicillin (HyClone), and 10 U mL^-1^ streptomycin (HyClone) at 37°C and 5% CO_2_. Cell passaging occurred upon reaching confluency in the flask by treating with 0.05% trypsin-EDTA for 4 minutes at 37°C and 5% CO_2_. Cells were pelleted at 500 *g* for 10 minutes, resuspended in 5 mL of complete media, and counted. HeLa cells were seeded in 12-well plates at 175,000 cells/well and 6-well plates at 750,000 cells/well. HEK293T cells were seeded in 6-well plates at 250,000 cells/well.

### Transfection

Plasmid DNA was prepared using ZymoPURE(tm) Plasmid Midiprep Kit (Zymo Research) according to the manufacture’s protocol. One day after seeding, transfection was achieved with Lipofectamine® 3000 (Invitrogen) using 1 μg total plasmid DNA per well for 6-well plates and 2.5 μg total plasmid DNA for 12-well plates in complete media and following the manufacture’s protocol. Where more than one plasmid was transfected, the total DNA was split evenly among all plasmids unless otherwise noted.

### Fluorescent microscopy and image analysis

All images were captured using an Observer Z.1 Inverted Microscope (Zeiss) with GFP, mCherry, BFP, or Cy5 filter cube sets (Chroma). For image analysis, five images were captured in each well. Image analysis was conducted using the “Measure” analysis in ImageJ with threshold set 10-255. Error bars on all plots represent the 95% confidence interval.

### Western blotting

Following imaging, cells were incubated in ice-cold lysis buffer (50 mM Tris, 150 mM NaCl, 1% Triton X-100, pH 8.0) on ice for 20 minutes with protease inhibitor cocktail (Calbiochem). Cells were then removed from the plate with a cell scrapper (Genemate), and the lysate was clarified in a pre-cooled centrifuge at 12,000 rpm for 10 minutes at 4°C. Total protein concentrations were normalized through a Bradford assay (Bio-Rad) with a BSA standard. 15 μg of lysate was mixed with a 5x loading buffer and separated by 10% SDS-PAGE before being transferred to a nitrocellulose membrane (Bio-Rad).

Western blots were blocked in TBST (20 mM Tris, 500 mM NaCl, 0.05% Tween-20, pH 8.0) containing 5% non-fat milk overnight at room temperature with gentle shaking. Membranes were washed twice in TBST and incubated for 3 hours in anti-GFP (1:5000 dilution, Covance) or anti-mCherry (1:2000 dilution, Novus) in TBS. The blots were then washed twice in TBST and incubated with horseradish peroxidase (HRP)-conjugated secondary antibody (GenScript) for 2 hours in TBST. The blots were washed three times in TBST and developed using ECL reagents (GE) according to the manufactures protocol. Band intensities were quantified using ImageJ gel analysis tools.

### Flow cytometry

Most flow cytometry was conducted on the Novocyte Benchtop Flow Cytometer (Acea Biosciences, San Diego, CA). Experiments involving mCherry (Ub R-nE7-mCherry and Ub X-LaM4-EGFP rescued with mCherry) were conducted on BD FACSAria Fusion High Speed Cell Sorter (BD Biosciences, San Jose, CA). All flow cytometry experiments involved ≥50,000 transfected cells as determined by forward- and side-scatter profiles of recorded events and fluorescent gating to exclude cells not transfected by at least the rescuing protein for each respective experiment. Cells were prepared for flow cytometry by washing twice in warm PBS. Trypsin treatment was applied for 3 minutes, and the reaction was quenched by warm media. Cells were collected in microcentrifuge tubes and spun at 0.8g for 5 minutes. The supernatant was aspirated, and cells were resuspended in cold PBS. This solution was then passed through a cell strainer into a flow cytometer tube and stored on ice until analysis.

### yCD viability studies

HeLa cells were seeded as above in 6-well plates and transfected with the appropriate constructs as above. Approximately one day post-transfection, wells either received no treatment, 5-FC, or 5-FU for 48 hours. Viability was determined using NucRed Dead 647 ReadyProbes Reagent (Thermo Fisher) per the manufacturer’s instruction. Fluorescent microscopy was used for analysis as described above.

### E7 detection studies

HeLa cells and HEK293T cells were seeded as described above. Transfection was conducted for 6 hours, and then replaced with normal media. Flow cytometry analysis was conducted 24 hours post-transfection as described above.

### Statistical analysis

All the experiments were performed in triplicates and results were expressed as means ± standard deviations (SD). Statistical significance was analyzed using the student *t*-tests. *P* < 0.05 was considered statistically significant throughout the study.

## Acknowledgments

This work was funded by grants from NSF (CBET1803008 and CBET1510817).

**The authors declare no conflict of interest.**

