## supplemental figures for "Conditional Protein Rescue (CPR) by Binding-Induced Protective Shielding"

**Supplementary Information Contents**

**Supplementary Figures**

**Supplementary Table**

**Supplementary Figures**

**1.	Fluorescent microscopy of SpyCatcher/SpyTag time course**

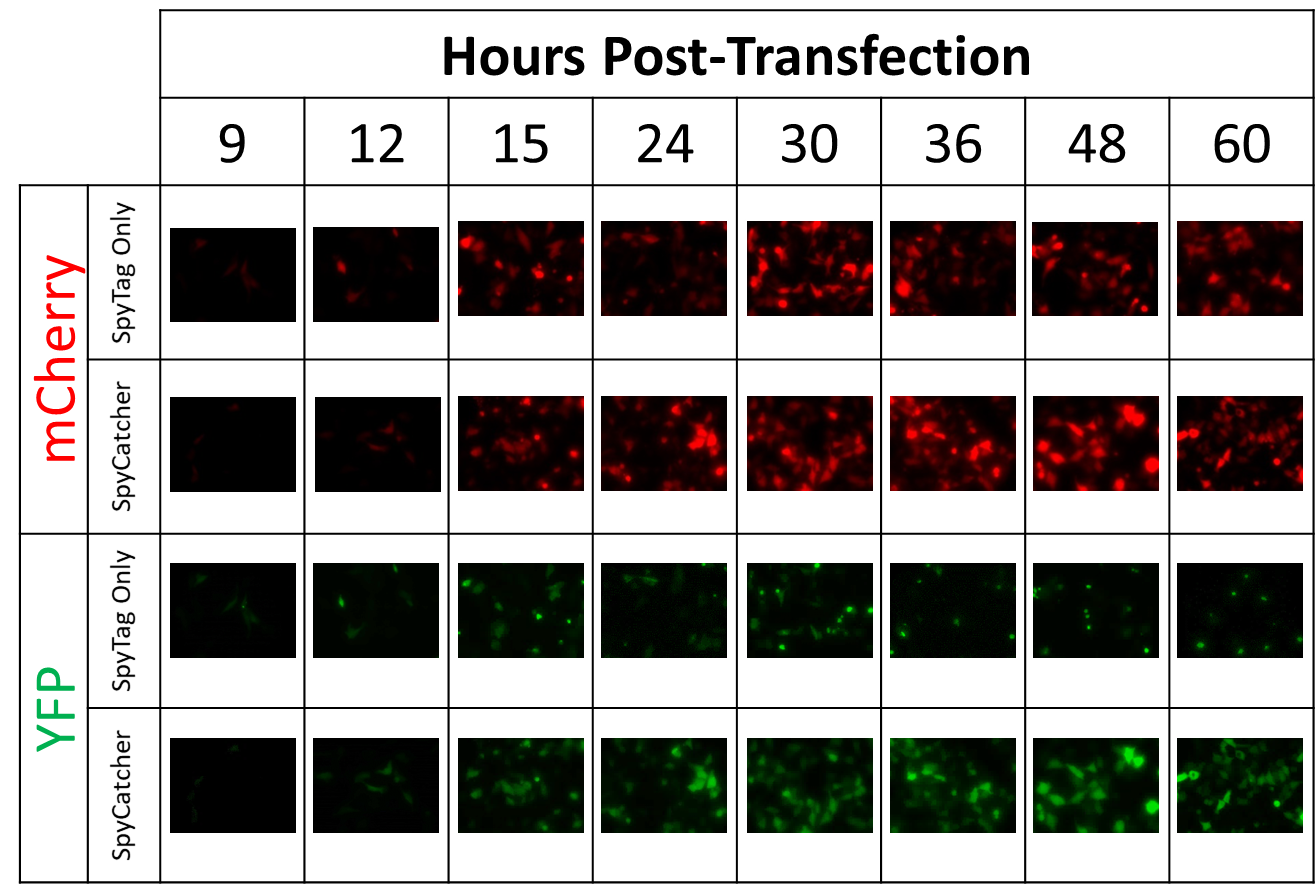

**Supplementary Figure 1.** **Fluorescent images of YFP rescue in the presence or absence of SpyCatcher-SpyTag ligation.** Representative images of HeLa cells transfected with mCherry T2A YFP-cODC1-SpyTag (SpyTag Only) or SpyCatcher-mCherry T2A YFP-cODC1-SpyTag (SpyCatcher) were captured over a 60-hour time course. While mCherry expression is consistent regardless of the presence of SpyCatcher, YFP expression is many-fold higher when SpyCatcher is co-expressed, suggesting that concealing the DD hinders the degradation of YFP.

**2.	Western Blot Validation of Flow Cytometry**

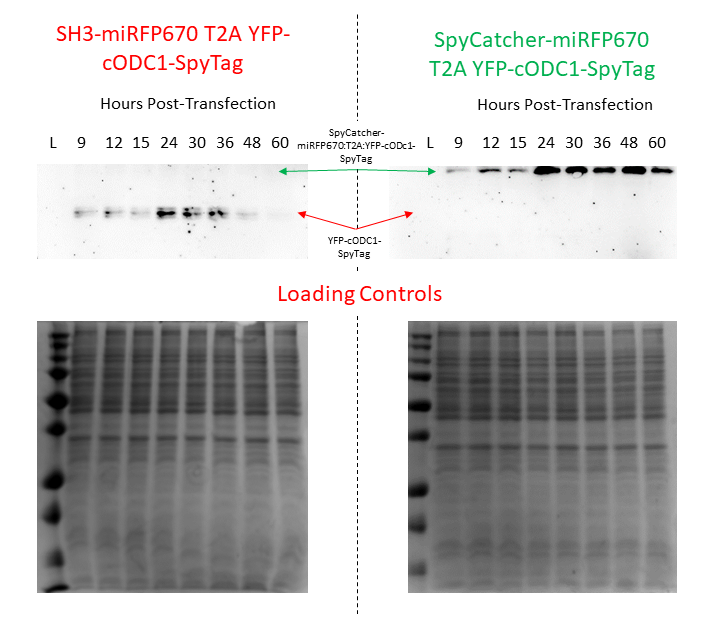

**Supplementary Figure 2.** **Western blot analysis demonstrating the rescue of YFP by ligating to SpyCatcher-miRFP670.** Substituting miRFP670 for mCherry in order to perform flow cytometry analysis has no effect on the level of YFP rescue. As with mCherry, the YFP signal degrades over time in the absence of SpyCatcher. When SpyCatcher is fused to mRFP670, there is a shift up in protein size indicative of SpyCatcher/SpyTag reaction, and the YFP signal persists strongly throughout the time course. The YFP signal increase is also reflected in the flow cytometry data in Fig. 2b.

**3.	Rescue by non-covalent interactions**

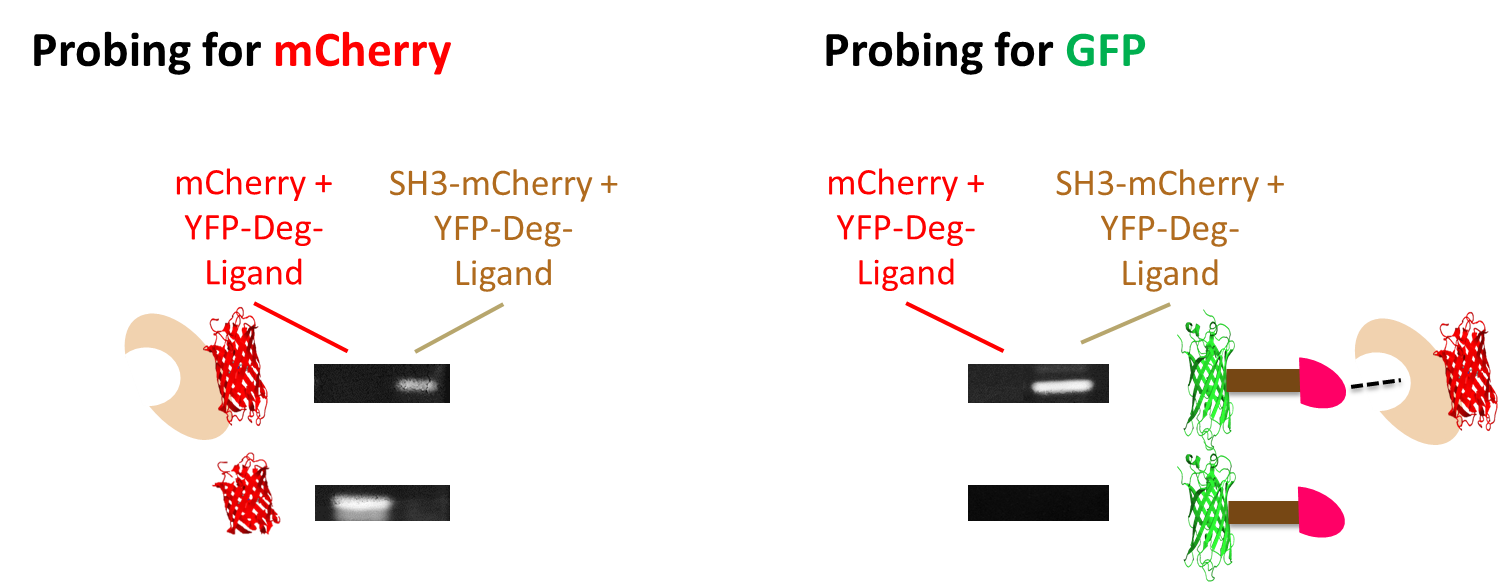

**Supplementary Figure 3.** **Non-covalent interactions resulting in protein rescue.** Western blots were generated after expressing mCherry T2A YFP-cODC1-Ligand (left lane in each pair) and SH3-mCherry T2A YFP-cODC1-Ligand (right lane in each pair) for 60 hours. These blots demonstrate the ability to rescue protein from degradation using an exogenous, non-covalent interacting pair. In this instance, the interacting protein pair is the SH3 domain and one of its known peptide ligands. When mCherry is expressed without an SH3 fusion, YFP is degraded by the degradation tag cODC1 (brown line). The close association between SH3 and its ligand is sufficient to induce YFP rescue by concealing the degradation tag from the proteasome.

**4.	GFP nanobody-mediated rescue**

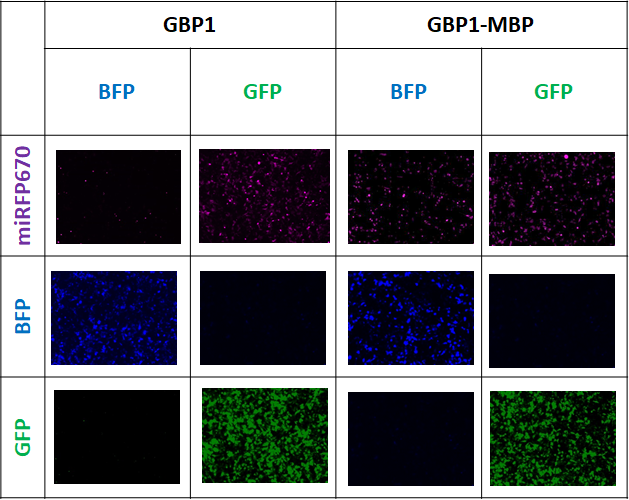

**Supplementary Figure 4. The use of nanobody-antigen interaction as an efficient sensing mechanism to control protein rescue.** Representative images of HeLa cells expressing either miRFP670-cODC1-GBP1 (left) or miRFP670-cODC1-GBP1-MBP (right) after 60 hours of transfection. The level of miRFP670 rescue was compared when either BFP or GFP was co-expressed at the same time. The miRFP670-cODC1-GBP1 sample receiving BFP shows low levels of miRFP670 due to its degradation. Both samples receiving GFP show rescue of miRFP670 from degradation, suggesting that the interaction between GFP and its nanobody concealed the degradation tag and rescued miRFP670. The direct fusion of MBP also stabilized miRFP670 even in the absence of GFP, suggesting a size-dependent steric interference model for CPR.

**5.	Effects of Rescuing Protein Orientation**

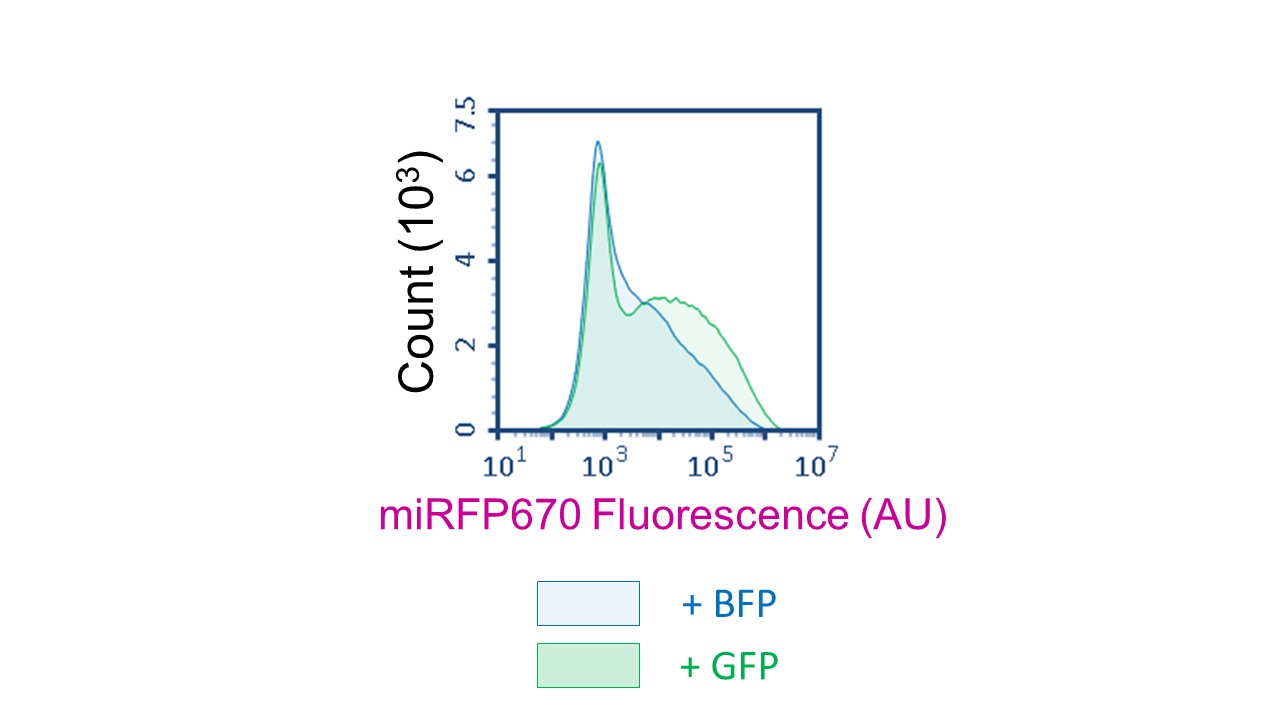

**Supplementary Figure 5. GBP6 fails to induce protein rescue effectively.** When GBP6, a nanobody that binds GFP at a different epitope than GBP1 (see Fig. 3 in the main text), is used to rescue miRFP670-cODC1-GBP6, the rescue efficiency is extremely low. This implies that GFP must interact at an orientation conducive to CPR in order to rescue miRFP670 from cODC1-mediated degradation.

**6.	UbL-mediated kinetics**

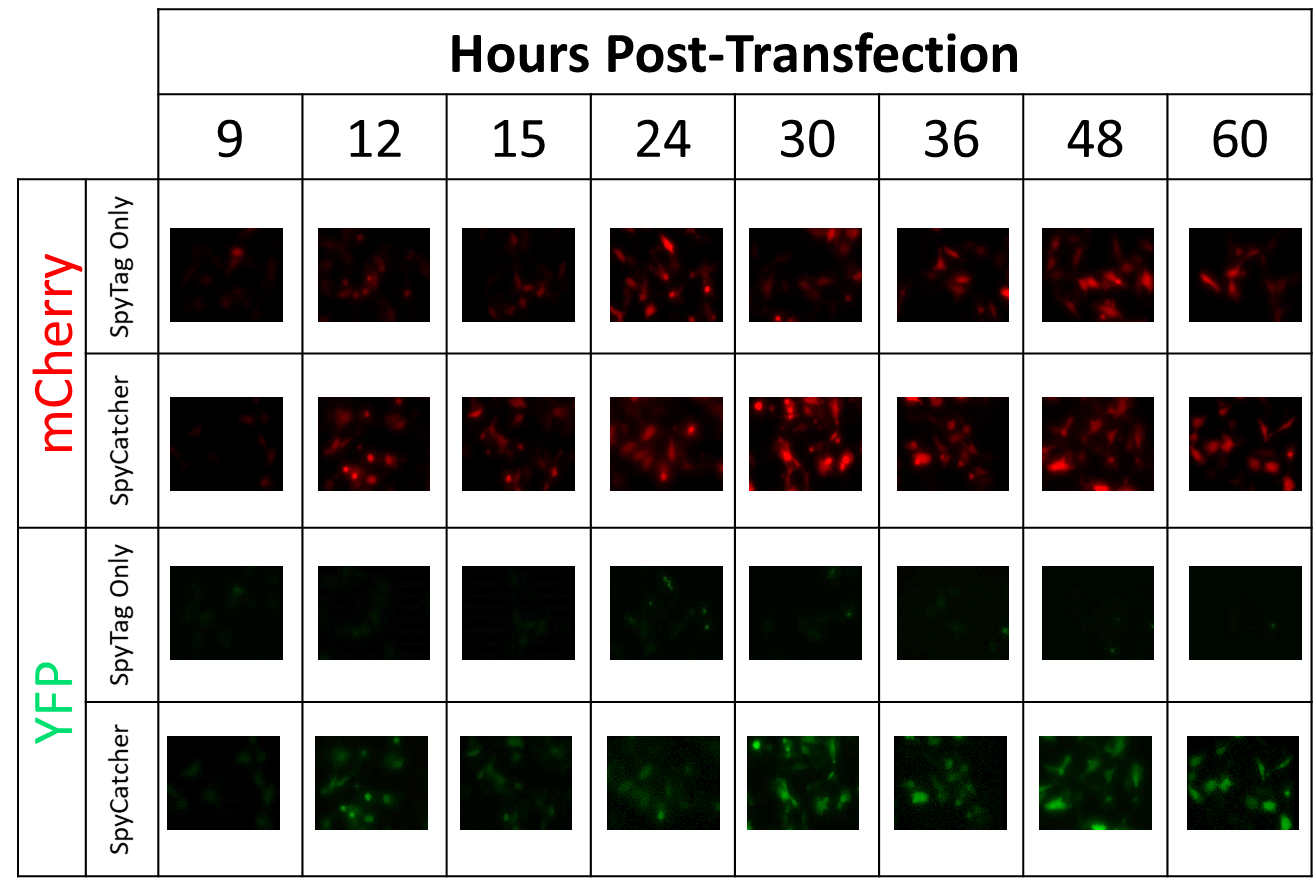

**Supplementary Figure 6. UbL-fusion speeds the degradation kinetics, resulting in lower background with efficient rescue.** Representative images of HeLa cells transfected with mCherry T2A UbL-YFP-cODC1-SpyTag (SpyTag Only) or SpyCatcher-mCherry T2A UbL-YFP-cODC1-SpyTag (SpyCatcher) were captured over a 60-hour time course. While overall YFP levels are lower when rescued compared to when UbL is not fused to its N-terminus, the background when SpyCatcher is not co-expressed is virtually non-existent.

**7.	N-End Rule-Mediated CPR of miRFP670**

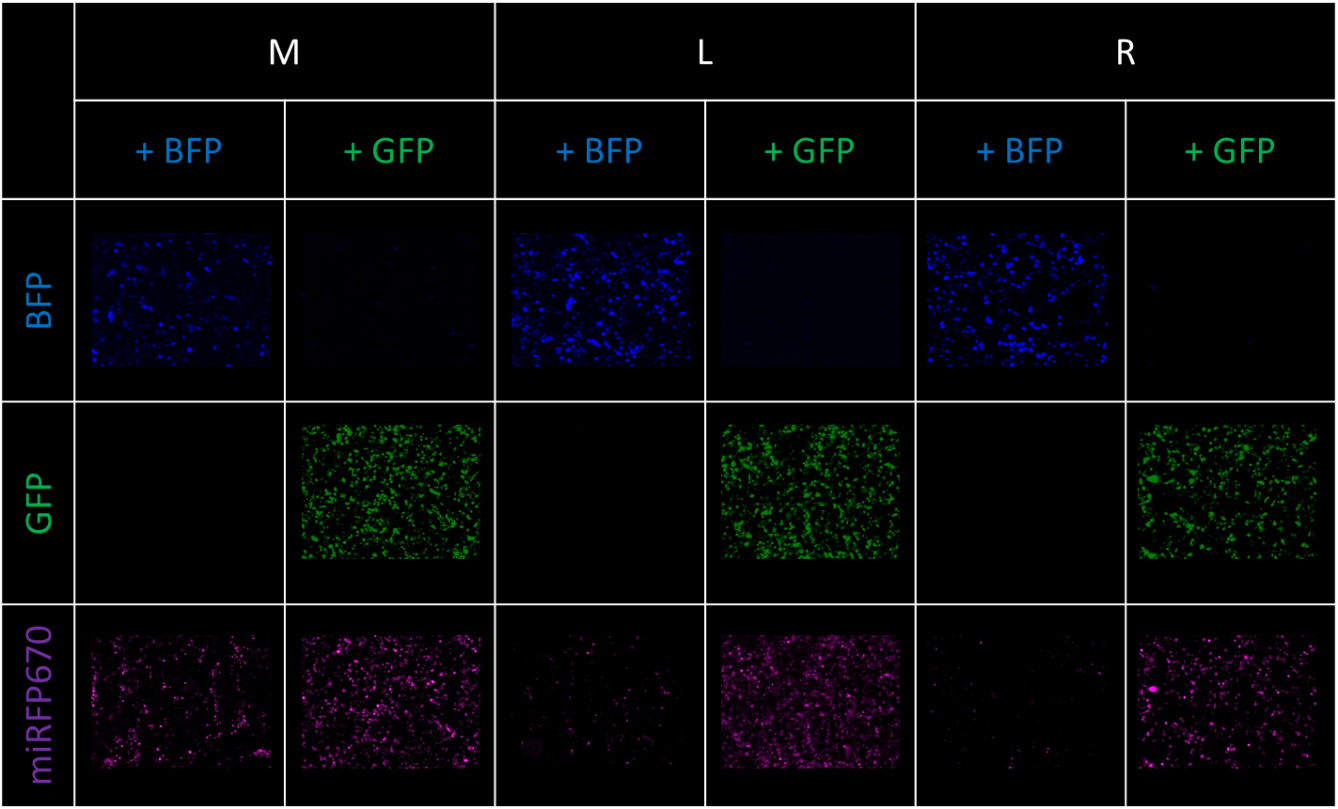

**Supplementary Figure 7. GFP is able to rescue miRFP670 tagged with degradation-inducing N-terminal amino acids by interacting with a GBP1.** Representative images of HEK293T cells transfected with X-GBP1-miRFP670, where X represents the N-terminal amino acids listed above. Co-expression with BFP resulted in low protein levels except when Met is the N-terminal amino acid, consistent with reported half-lives. Co-expression with GFP results in a visible increase in miRFP670 in the case of all N-terminal amino acids.

**8.	CPR Operates with Different Nanobodies and Target Proteins**

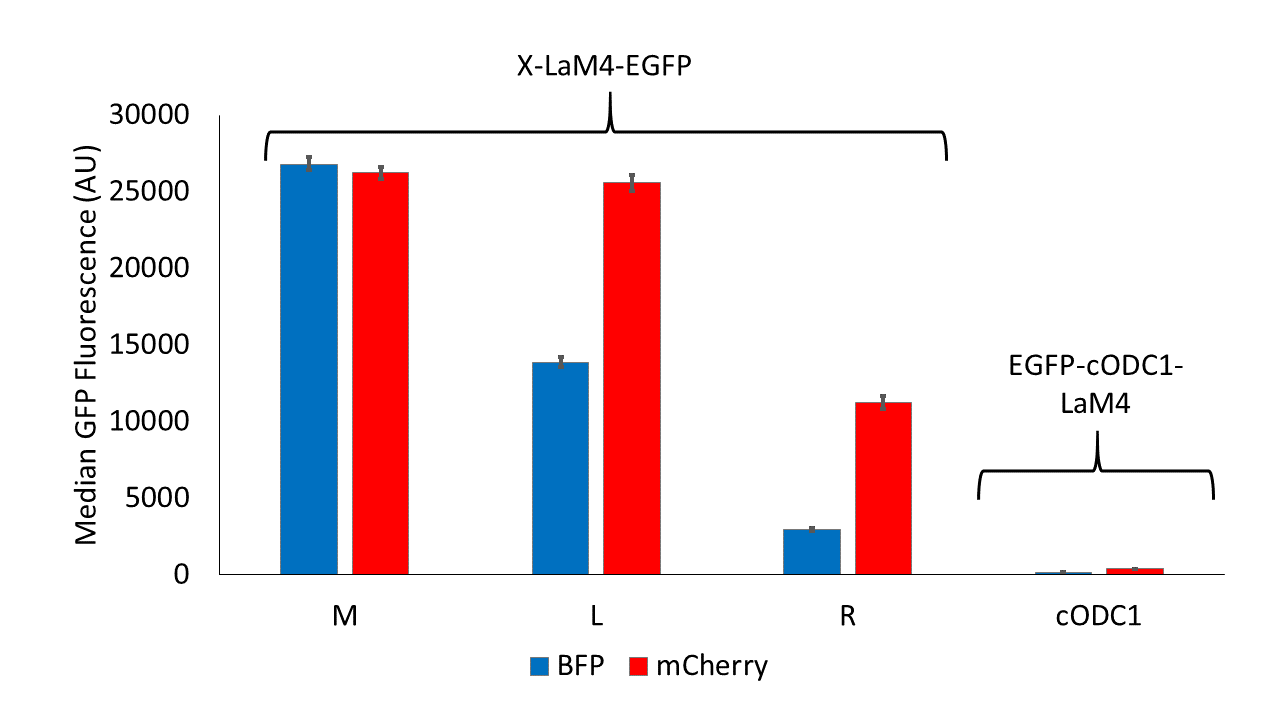

**Supplementary Figure 8. mCherry is able to rescue EGFP by interacting with a LaM4.** When co-expressed with BFP, levels of EGFP scale well with the reported half-lives of the N-terminal amino acids, and EGFP-cODC1-LaM4 exhibits low levels of EGFP fluorescence. However, co-expression with mCherry results in levels of EGFP in all samples. Results graphed represent the median fluorescence values measured by flow cytometry, and error bars represent 95% confidence intervals.

**Supplementary Tables**

**Supplementary Table 1.** All oligonucleotides were ordered from Integrated DNA Technology (Coralville, IA), purified via standard desalting, and ordered as a standard oligo unless otherwise noted.

| **Name** | **Category** | **Sequence (5’🡪3’)** |
| --- | --- | --- |
| *AflII* YFP | Forward PCR Primer | atatatttaaGatgGTGAGCAAGGGCGAGGA |
| YFP *XhoI* | Reverse PCR Primer | atatatctcgagcttgtacagctcgtccatg |
| *XhoI* cODC1-SpyTag *ApaI* | Ultramer (Sense) | tcgagATGTCTTGTGCCCAAGAGTCAATAACCAGTCTGTATAAGAAAGCTGGAAGTGAAAACCTCTATTTTCAGtctagAGCACACATAGTAATGGTAGACGCCTACAAGCCGACGAAGtaaGggcc |
|  | Ultramer (Antisense) | CttaCTTCGTCGGCTTGTAGGCGTCTACCATTACTATGTGTGCTctagaCTGAAAATAGAGGTTTTCACTTCCAGCTTTCTTATACAGACTGGTTATTGACTCTTGGGCACAAGACATc |
| *NheI* mCherry | Forward PCR Primer | ATATATgctagcATATTAgctaagcATGGTGAGCAAGGGCGAGGA |
| T2A mChery *AflII* | Reverse PCR Primer | atatatCTTAAGggggccggggttctcctccacgtcgccgcaggtcagcagggagcccctgccctcATTCTTGTACAGCTCGTCCA |
| Human Codon Optimized SpyCatcher | gBlock Gene Fragment | GACAGCGCCACCCACATCAAGTTCAGCAAGAGGGACGAGGACGGCAAGGAGCTGGCCGGCGCCACAATGGAGCTGAGAGACAGCAGCGGCAAGACCATCAGCACCTGGATCAGCGACGGCCAGGTGAAGGACTTCTACCTGTACCCCGGCAAGTACACCTTCGTGGAGACCGCCGCCCCCGACGGCTACGAGGTGGCCACCGCCATCACCTTCACCGTGAACGAGCAGGGCCAGGTGACCGTGAACGGC |
| *NheI* SpyCatcher | PCR Forward Primer | atatatgctagcATGGACAGCGCCACCCACATCAAGTTCA |
| SpyCatcher *EcoRI* | PCR Reverse Primer | atatatgaattcGCCGTTCACGGTCACCTGGCCCTGC |
| *EcoRI* mCherry | PCR Forward Primer | atatatgaattcATGGTGAGCAAGGGCGAGGA |
| *NheI* miRFP670 | PCR Forward Primer | atatatgctagccgccaccATGGTGGCTGGACACGCTTC |
| miRFP670 *HindIII* | PCR Reverse Primer | atatataagcttGCTCTCCAGGGCGGTGATTC |
| *HindIII* T2A *AflII* | Oligo (Sense) | agcttgagggcaggggctccctgctgacctgcggcgacgtggaggagaaccccggccccc |
|  | Oligo (Antisense) | ttaagggggccggggttctcctccacgtcgccgcaggtcagcagggagcccctgccctca |
| *EcoRI* miRFP670 | PCR Forward Primer | atatatgaattcATGGTGGCTGGACACGCTTC |
| *NheI* SH3 | PCR Forward Primer | atatatgctagcATGGCAGAGTATGTGCGGgc |
| SH3 *EcoRI* | PCR Reverse Primer | atatatgaattcATACTTCTCCACGTAAGGGA |
| *XbaI* SH3Lig *ApaI* | Oligo (Sense) | ctagaCCACCACCAGTCCCCCCTAGACGAtaagggcc |
|  | Oligo (Antisense) | tGGTGGTGGTCAGGGGGGATCTGCTattc |
| *XbaI* SpyTag *BamHI* | Oligo (Sense) | ctagaGCACACATAGTAATGGTAGACGCCTACAAGCCGACGAAGg |
|  | Oligo (Antisense) | gatccCTTCGTCGGCTTGTAGGCGTCTACCATTACTATGTGTGCt |
| *BamHI* GBP1 | PCR Forward Primer | atatatggatccATGCAGGTGCAACTGGTGGA |
| GBP1 *ApaI* | PCR Reverse Primer | atatatgggcccttaCTTGCTGCTCACGGTCACCTGGGTG |
| *AflII* miRFP670 | PCR Forward Primer | atatatcttaagATGGTGGCTGGACACGCTTC |
| miRFP670 *XhoI* | PCR Reverse Primer | atatatctcgagGCTCTCCAGGGCGGTGATTC |
| *NheI* BFP | PCR Forward Primer | atatatGCTAGCataaGAATTCatgagcgagctgattaaggagaacatgcaca |
| BFP *ClaI* | PCR Reverse Primer | atatatatcgatGTTCAGCTTgtgccccagtttgctagggaggtc |
| *ClaI* T2A *AflII* | Ultramer (Sense) | cgatGGCAGCGGCGAGGGCAGAGGCAGCCTGCTGACCTGCGGCGACGTGGAGGAGAACCCCGGCCCCc |
|  | Ultramer (Antisense) | ttaagGGGGCCGGGGTTCTCCTCCACGTCGCCGCAGGTCAGCAGGCTGCCTCTGCCCTCGCCGCTGCCat |
| *EcoRI* BFP | PCR Forward Primer | atatatgaattCatgagcgagctgattaagga |
| *AflII* yCD | PCR Forward Primer | atatatcttaagatgGTGACCGGCGGAATGGCCAG |
| yCD *XhoI* | PCR Reverse Primer | atatatctcgagGCTGCCggatccTTCTCCAATATCCTCAAACC |
| *NheI* EGFP | PCR Forward Primer | atatagctagcATGGTGAGCAAGGGCGAGGagctgt |
| EGFP *EcoRI* | PCR Reverse Primer | atatatgaattcGCTGCCGCCGCCCTTGTACAGCTCGTCCATGCCgaga |
| *AflII* UbL | PCR Forward Primer | atatatcttaagATGCAGGTGACCCTGAAGACCCTGC |
| UbL *ClaI* | PCR Reverse Primer | atatatatcgatAGAACCACCACCAGATCCACCGTCT |
| *ClaI* YFP | PCR Forward Primer | atatatatcgatGTGAGCAAGGGCGAGGAgctgttca |
| Ub-M-EGFP | Mutagenesis Forward Primer | CAGAGGTGGGATGGGGAAGCTTGGTCG |
| Ub-L-EGFP | Mutagenesis Forward Primer | CAGAGGTGGGCTGGGGAAGCTTG |
| Ub-X-EGFP | Mutagenesis Reverse Primer | AGACGGAGTACCAGGTGC |
| Ub-X-KpnI_BamHI-EGFP | Mutagenesis Forward Primer | ATATGGATCCATGGTGAGCAAGGGCG |
|  | Mutagenesis Reverse Primer | ATGGTACCTTGTCGACCAAGCTTCCC |
| *KpnI* GBP1 | PCR Forward Primer | atatatggtaccATGCAGGTGCAACTGGTGGA |
| GBP1 *BamHI* | PCR Reverse Primer | atatatggatccCTTGCTGCTCACGGTCACCT |
| *BamHI* miRFP670 | PCR Forward Primer | atatatGGATCCATGGTGGCTGGACACGCTTCCGG |
| miRFP670 *NotI* | PCR Reverse Primer | atatatgcggccgcttaGCTCTCCAGGGCGGTGAT |
| *AflII* EGFP | PCR Forward Primer | atatatcttaagATGGTGAGCAAGGGCGAGGa |
| EGFP *XhoI* | PCR Reverse Primer | atatatctcgagCTTGTACAGCTCGTCCATGC |
| *BamHI* LaM4 | PCR Forward Primer | atatatggatccATGGCTCAGGTGCAGCTCGTGGA |
| LaM4 *ApaI* | PCR Reverse Primer | atatatgggcccttaGGTGAAAGGGGAAGACACGG |
| *KpnI* LaM4 | PCR Forward Primer | atatatggtaccATGGCTCAGGTGCAGCTCGT |
| LaM4 *BamHI* | PCR Reverse Primer | atatatggatccGGTGAAAGGGGAAGACACGG |
| *KpnI* nE7 | PCR Forward Primer | atatatggtaccatgCAGGTTCAGCTGGTGGAAAG |
| nE7 *BamHI* | PCR Reverse Primer | atatatggatccGCTGCTCACGGTCACCTGGG |
| *AgeI* mCherry | PCR Forward Primer | atatataccggtcgccaccATGGTGAGCAAGGGCGAGGA |
| mCherry *NotI* | PCR Reverse Primer | atatatcgcggccgctttaCTTGTACAGCTCGTCCATGC |
